# Protein-Independent Liquid–Liquid Phase Separation of Adenosine Triphosphate under Crowded Conditions

**DOI:** 10.1101/2025.11.24.690196

**Authors:** Robert Dec, Wojciech Dzwolak, Roland Winter

**Affiliations:** Physical Chemistry I - Biophysical Chemistry, Department of Chemistry and Chemical Biology, TU Dortmund University, Otto-Hahn Strasse 4a, 44227 Dortmund, Germany; Faculty of Chemistry, Biological and Chemical Research Centre, University of Warsaw, Pasteur Street 1, 02-093 Warsaw, Poland

## Abstract

Adenosine triphosphate (ATP), a common constituent of protein-rich biomolecular condensates, itself undergoes liquid–liquid phase separation (LLPS) even in the absence of polypeptides. Using polyethylene glycol as macromolecular crowding agent, we observed robust ATP droplet formation at pH 2–11. Moderate NaCl and lower temperatures promote LLPS by lowering critical ATP and crowder concentrations. Very high salt reverses this trend through anomalous underscreening, but ATP droplets still survive in hypersaline environments. Most importantly, physiological millimolar ATP concentrations are sufficient for phase separation in the presence of millimolar Mg^2+^ and crowders, mimicking intracellular conditions. pH tunes intermolecular interactions, evidenced by inversion of the adenosine circular dichroism Cotton effect. These results reveal intrinsic, protein-independent LLPS of ATP with potential roles in cellular compartmentalization and pathological phase transitions.

---

Liquid–liquid phase separation (LLPS) is a ubiquitous physicochemical process (1) that drives the assembly of membraneless organelles such as nucleoli, P bodies, and stress granules (2-7). It has also been proposed that LLPS enabled the emergence of prebiotic protocells by furnishing a primitive yet effective route to compartmentalization (8–11). Conversely, dysregulated protein phase separation is increasingly associated with the early stages of pathological aggregation in neurodegenerative diseases and beyond, such as by α-synuclein, FUS, tau and PrP (12-20). Adenosine triphosphate (ATP), best known as the universal energy currency of the cell, also exerts profound effects on biomolecular condensation. The molecule exhibits an amphiphilic nature due to the presence of both charged, hydrophilic groups, the triphosphate, and a hydrophobic adenosine ring system. At millimolar physiological concentrations, ATP can either promote or suppress protein phase separation, depending on the system. While the mechanisms by which ATP affects proteins in solution are nuanced and still under debate, the nucleotide often acts as a potent hydrotrope that solubilizes hydrophobic polypeptides, inhibits amyloid formation, and dissolves pre-existing aggregates, but may also promote aggregation of neutral macromolecules, i.e., show salting-out behavior (21-24). In other contexts, ATP actively drives LLPS by bridging multivalent cationic proteins and peptides through electrostatic and cation– π interactions, as observed with FUS (25, 26), arginine-(27) or lysine-rich (28-31) peptides, IgG1 antibodies (32), and the N-protein of SARS-CoV-2 (33). Depending on its concentration, ATP can promote either type of transition (25, 34). In all cases reported to date, however, ATP acts as a minor component or modulator within protein- or peptide-rich condensates.

Here, we report the unexpected discovery that ATP itself—without any protein, peptide, or nucleic acid—undergoes robust liquid–liquid phase separation under conditions of macromolecular crowding that mimic the dense intracellular environment. This intrinsic, protein-independent phase behavior of ATP reveals a hitherto unrecognized dimension of this quintessential metabolite and raises new questions about its potential roles in the formation of cellular condensates, their regulation, and pathology-associated phase transitions.

## Results and discussion

### Crowder-induced phase separation and droplet formation of ATP

Like many amphiphilic molecules in aqueous solution, ATP tends to self-associate at the molecular and microscopic levels or forms nanoclusters as revealed by NMR-monitoring (24, 35-40). However, the observation reported here of phase-separating liquid droplets formed by ATP in the absence of proteinaceous partners under molecular crowding conditions (known to favor droplet formation (41-44)) emerged unexpectedly. Routine microscopic inspection of mixed aqueous ATP-PEG samples used as control in a different project revealed the presence of highly dynamic droplets (Fig. 1A, Supporting Movies 1 and 2). The droplets formed rapidly in a very broad pH range (from 2 to 11) as long as the concentrations of both ATP and PEG were above certain critical levels. Optical density (turbidity) measurements at 500 nm were selected as a high-throughput means of quantifying the abundance of droplets. The corresponding plots are shown in Figs. 1B and 1C. In concentrated (33.3 % wt./vol.) PEG-3350 solution at 21 °C, LLPS occurred at ATP concentrations above 55 or 35 mg/mL, depending on whether the process took place at pH 7 or 4, clearly indicating that the protonation-induced decrease in repulsion between unscreened ATP ions favors droplet formation. The phase separation showed an upper critical solution temperature (UCST) phase behavior: raising the temperature from 21 °C led to mixing, while cooling the samples prepared at subcritical concentrations triggered demixing below ∼10 °C (Figs. 1D, E).

**Fig. 1.**
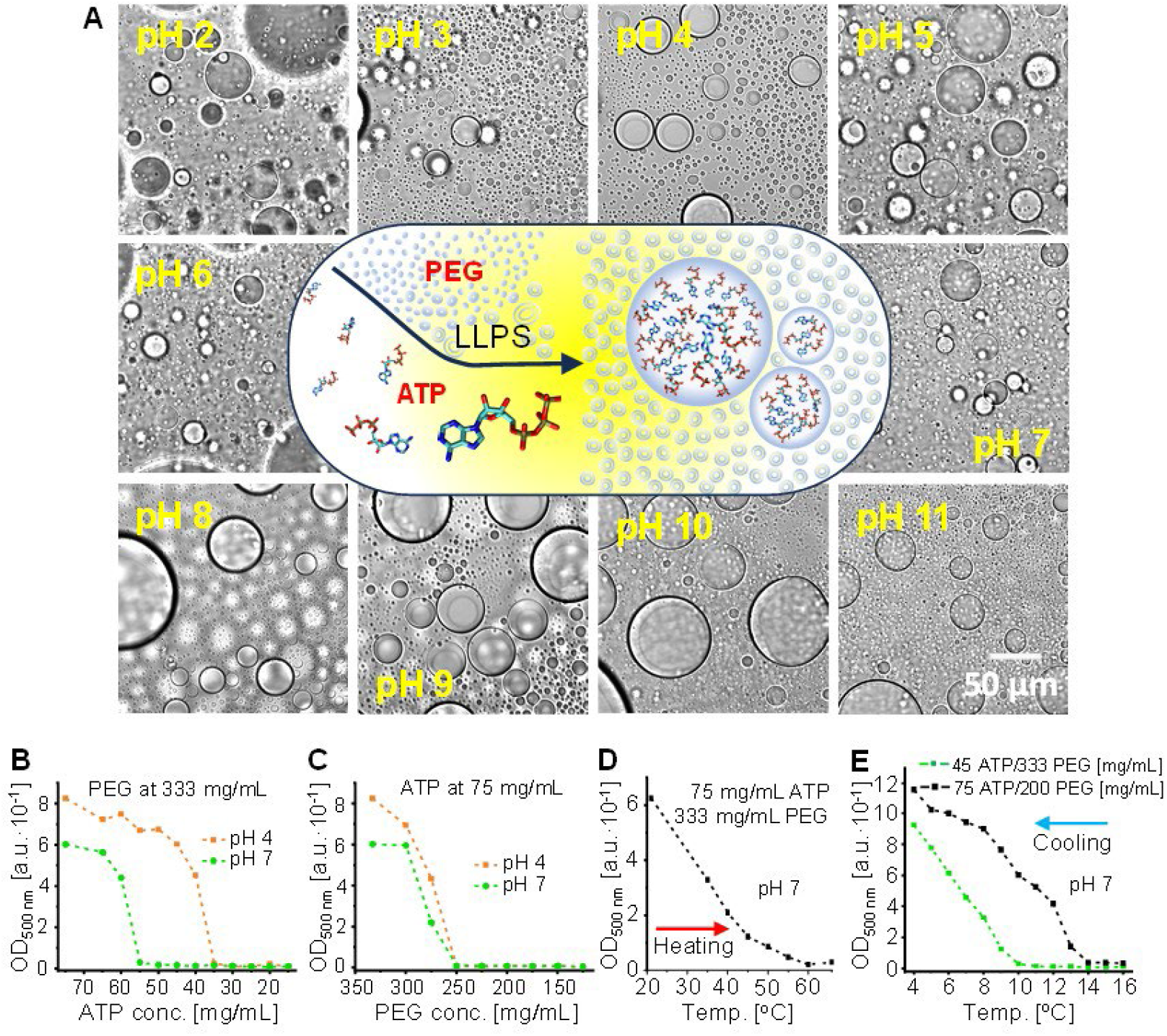
Under molecular crowding conditions, ATP forms liquid droplets on its own in a broad pH range. Brightfield microscopic images of droplets formed in aqueous solutions containing 75 mg/mL ATP and 333 mg/mL PEG-3350 (average MW of 3350 Da) with pH adjusted to the indicated values; all images were acquired at room temperature (A). Optical density (absorbance at 500 nm) arising from the LLPS as a function of ATP concentration (at fixed 333 mg/mL PEG-3350 concentration - B) and of PEG-3350 concentration (at fixed 75 mg/mL ATP concentration - C) at pH 4 and 7 and at 21 °C. Temperature-induced mixing in the two-phase system formed in the presence of 75 mg/mL ATP and 333 mg/mL PEG-3350, pH 7, probed by absorbance measurements at 500 nm (D). Cooling-induced LLPS in the single-phase systems with initial subcritical concentrations of ATP (45 ATP/333 PEG [mg/mL]) and PEG-3350 (75 ATP/200 PEG [mg/mL]), pH 7 (E).

Under the various conditions investigated here, the ATP droplets coalesced rapidly (see Supporting Movies 1 and 2) and formed a macroscopically separated coacervate phase (Fig. 2A), which accounted to less than 10 % of the total volume (subject to the pH conditions and the exact mixing stoichiometry). The segregated system is quite stable even under ambient conditions (see Fig. S1). The different ATP concentrations in the upper and bottom liquid phases were revealed by the results of luciferase-based ATP activity assays. According to the spectrophotometric estimation of ATP concentrations at three different pH values (see inset Table in Fig. 2A), the nucleotide concentration in the bottom layer is at least one order of magnitude higher than in the upper phase and continues to increase as the pH decreases (∼320 mg/mL at pH 9 compared to ∼390 mg/mL at pH 7 and to ∼437 mg/mL at pH 4).

**Fig. 2.**
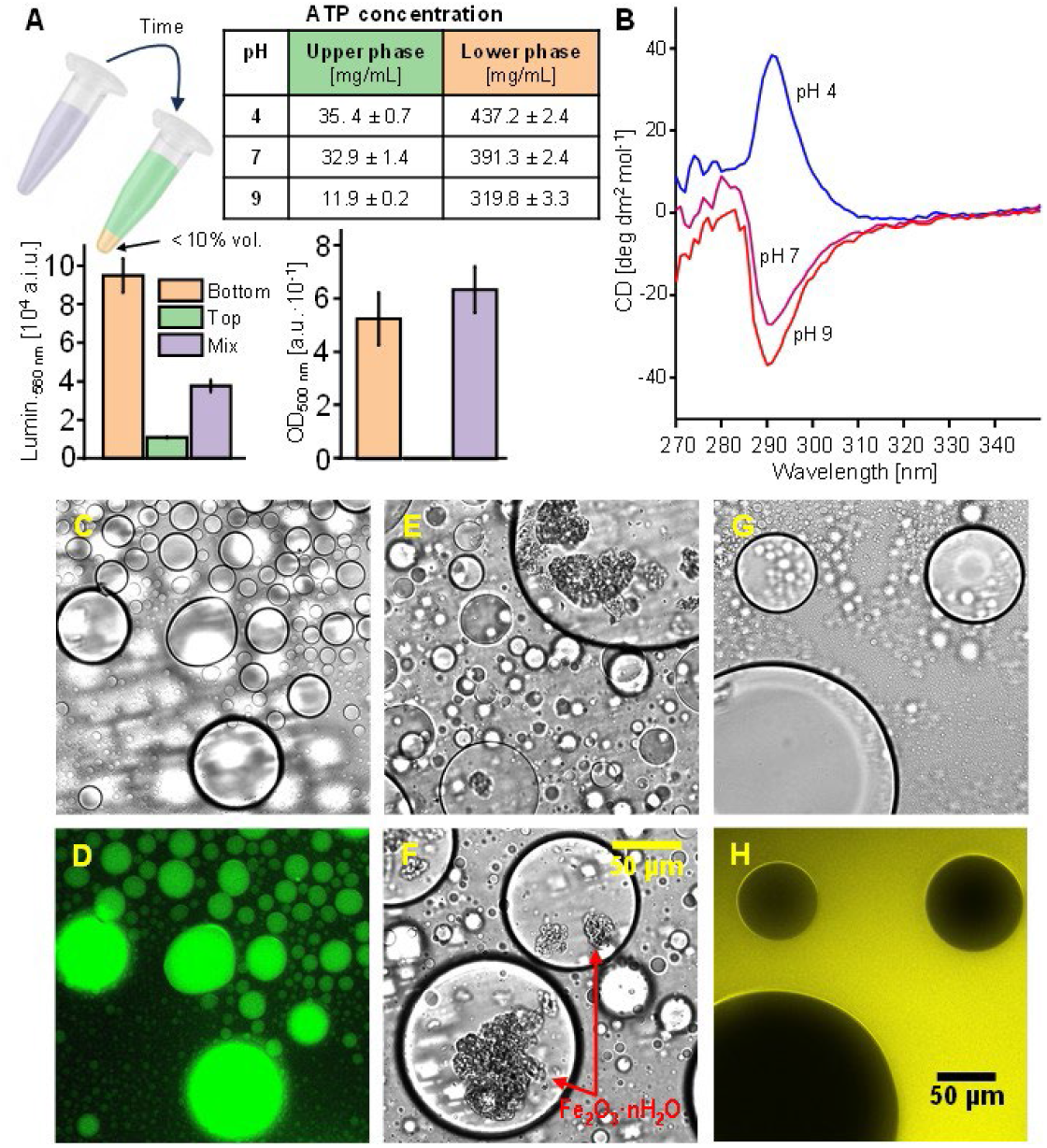
Composition and properties of the ATP droplets. Droplets formed in aqueous mixture of 75 mg/mL ATP and 333 mg/mL PEG-3350, pH 7, readily coalesce forming two macroscopic phases of which the bottom ATP-rich phase amounts to less than 10 % of the total volume (A). Relative luminescence levels (at 560 nm) from a luciferase-based ATP assay juxtaposed to corresponding optical densities (at 500 nm) of the top and bottom phases before and after separation; Table inset: concentrations of ATP in the ATP-rich and top phases at pH 4, 7, and 9, obtained from spectrophotometric measurements (at 260 nm) of water-diluted aliquots. CD spectra of centrifuged ATP-rich phases at pH 4, 7, and 9 (75 mg/mL ATP + 333 mg/mL PEG-3350, 25 °C), measured using a 0.01 mm optical pathway and subsequently normalized to the ATP concentration (B). Brightfield (C, E, G, F) and fluorescence (D, H) microscopy images of 75 mg/mL ATP, 333 mg/mL PEG-3350, pH 7 droplets containing 0.02 mg/mL quinacrine (C, D), microparticles of sonicated Fe_2_O_3_·nH_2_O (E, F) or 0.2 mg/mL fluorescent PEG-FITC (G, H). The fluorescence was excited at excitation 480/40 and at emission 527/30, both for PEG-FITC and quinacrine.

Circular dichroism (CD) spectra of the ATP-rich bottom phases at these three different pH values were collected using a quartz cuvette with an extremely short optical pathway of 0.01 mm (Fig. 2B). We noted that the sign of the Cotton effect induced in the adenine chromophore switches from negative to positive when the pH is lowered from 7 to 4, indicating distinct patterns of intermolecular association and hence transition dipole moment-couplings between the ATP molecules, which is reminiscent of a similar behavior of solution-phase ATP oligomers (36).

### Partitioning of cosolutes in ATP droplets

We have subsequently used brightfield and fluorescence microscopy to probe the droplets’ capacity to sequester selected foreign molecules and particles (Figs. 2C to 2H). While fluorescent quinacrine (Figs. 2C, 2D), and to some extent inorganic Fe_2_O_3_·nH_2_O microparticles (45) readily accumulate within the ATP droplets (Figs. 2E, 2F), fluorophore-labelled PEG does not enter the droplets, as shown in Figure 2H. Thus, PEG is completely excluded from the ATP-rich droplet phase.

### Salt-effects on phase separation and droplet formation of ATP - screening and underscreening

As the interplay of hydrophobic, π-stacking, and Coulombic interactions clearly affects the stability and dynamics of ATP droplets, we investigated the LLPS transition in the presence of moderately concentrated NaCl, which provides Debye screening for the repulsive forces between nucleotide polyanions. The optical density traces and microscopic images in Fig. 3 show how the presence of 2 M NaCl lowers the critical concentrations of ATP and PEG (panels A and B, respectively) at which LLPS occurs. The effect of salt at this concentration is profound: when all other parameters (except for ionic strength) are kept constant, the droplet state becomes accessible to ATP at concentrations approximately 4 times lower. Likewise, the critical concentration of PEG is reduced by about 40 % (Fig. 3B compared to Fig. 1C).

**Fig. 3.**
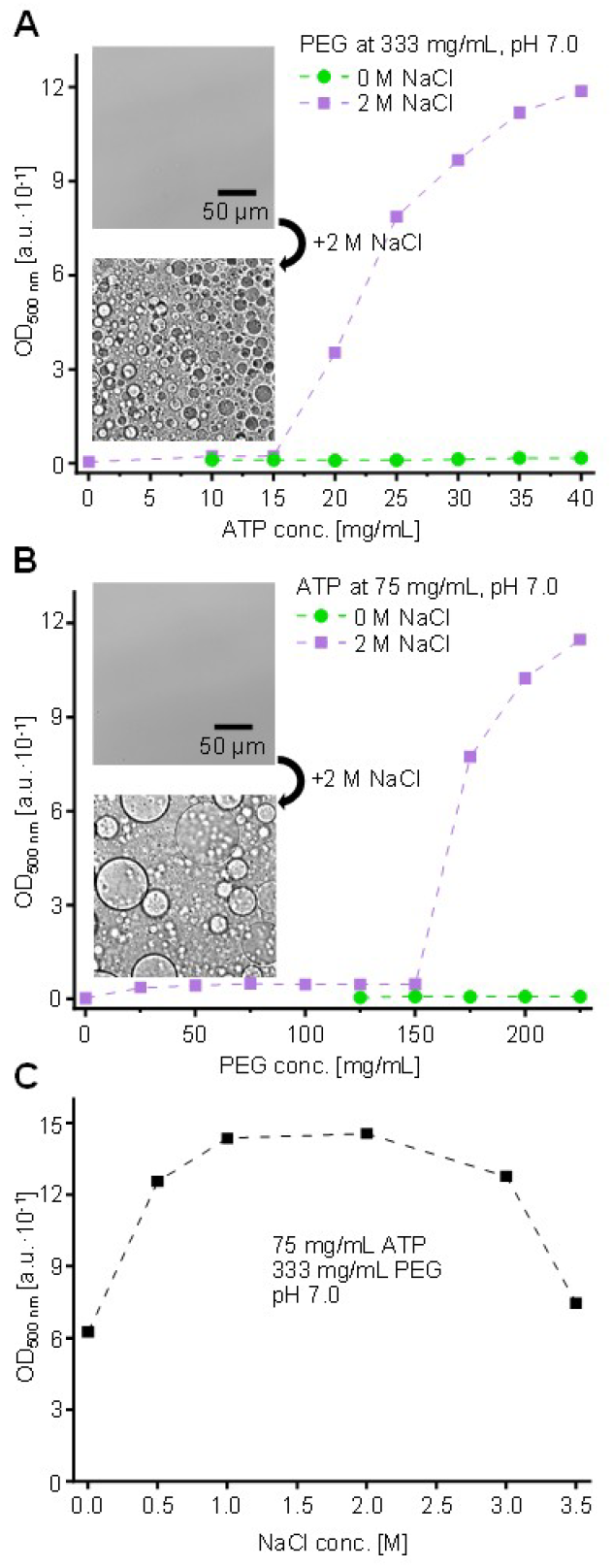
Impact of high ionic strength on LLPS in the ATP-PEG-3350 system. Optical density changes accompanying titration of PEG (333 mg/mL, pH 7) with ATP (A) and vice versa (75 mg/mL ATP, pH 7 - B) in the absence and presence of 2 M NaCl (both at 21 °C). Microscopic images of liquid 25 mg/mL ATP + 333 mg/mL PEG in the absence and presence of 2 M NaCl are presented in panel A, whereas the inset in panel B shows the analogous images collected for 75 mg/mL ATP + 175 mg/mL PEG (i.e., at subcritical for LLPS concentrations of ATP and PEG, respectively). Non-monotonic dependence of the optical density of the 75 mg/mL ATP + 333 mg/mL PEG, pH 7 system on NaCl concentration (C).

Interestingly, a more comprehensive investigation of the effect of NaCl concentration on LLPS in the ATP-PEG system (probed by turbidity measurements) presented in Fig. 3C suggests that Debye screening becomes less effective at extremely high salt concentrations (above 2 M NaCl). This puzzling effect can be attributed to the recently explored phenomenon of anomalous underscreening, which occurs when the mean electrostatic potential is less screened than predicted by Debye-Hückel theory and the electrostatic decay length becomes significantly larger than predicted by the classical mean-field approach (46-48). This phenomenon is associated with pronounced short-range forces (e.g., hydrophobic and π-driven interactions) that come into play, leading to cluster and ion pair formation (46-48). However, ATP droplets still exist at the highest NaCl concentration recorded (3.5 M). It is therefore noteworthy that ATP droplets could even survive hypersaline salt environments, such as those found on Earth or in the salt-lakes of the Martian subsurface (49), and that they could play a role in the development of prebiotic cells under such harsh stress conditions.

The data presented in Figure 4 indicate that millimolar concentrations of Mg^2+^ cations, which naturally complex with cellular ATP, make the LLPS transition even more accessible than non-physiologically concentrations of NaCl. Specifically, at 50 mM MgCl_2_, ambient temperature, and neutral pH, droplets begin to form at a concentration of 2.5 mg/mL ATP (∼ 5 mM), which is within the physiological range (49, 50) (see the sparse complex droplets in the left section of Fig. 4A). Of note, ATP concentrations can reach much higher values in specific cellular compartments, such as 0.1 M in the adrenal medulla (24). The increased droplet formation continues even when the Mg^2+^ concentration is reduced to 30 mM (Fig. 4B). Moreover, we observed that tiny dispersed droplets form near the glass-solution interfaces even when the PEG concentration is reduced to 17.5 % wt./vol. (at 30 mM MgCl_2_ and 7 mg/mL ATP, pH 7) and the temperature is reduced to 4 °C (Fig. 4C). One of the characteristic features of Mg^2+^-stabilized ATP droplets is their persistence in solution in the form of a finely dispersed emulsion that coalesces into the macroscopically separating phase over much longer periods of time than the droplets formed in the absence of Mg^2+^ and at high nucleotide concentration (Fig. 4D).

**Fig. 4.**
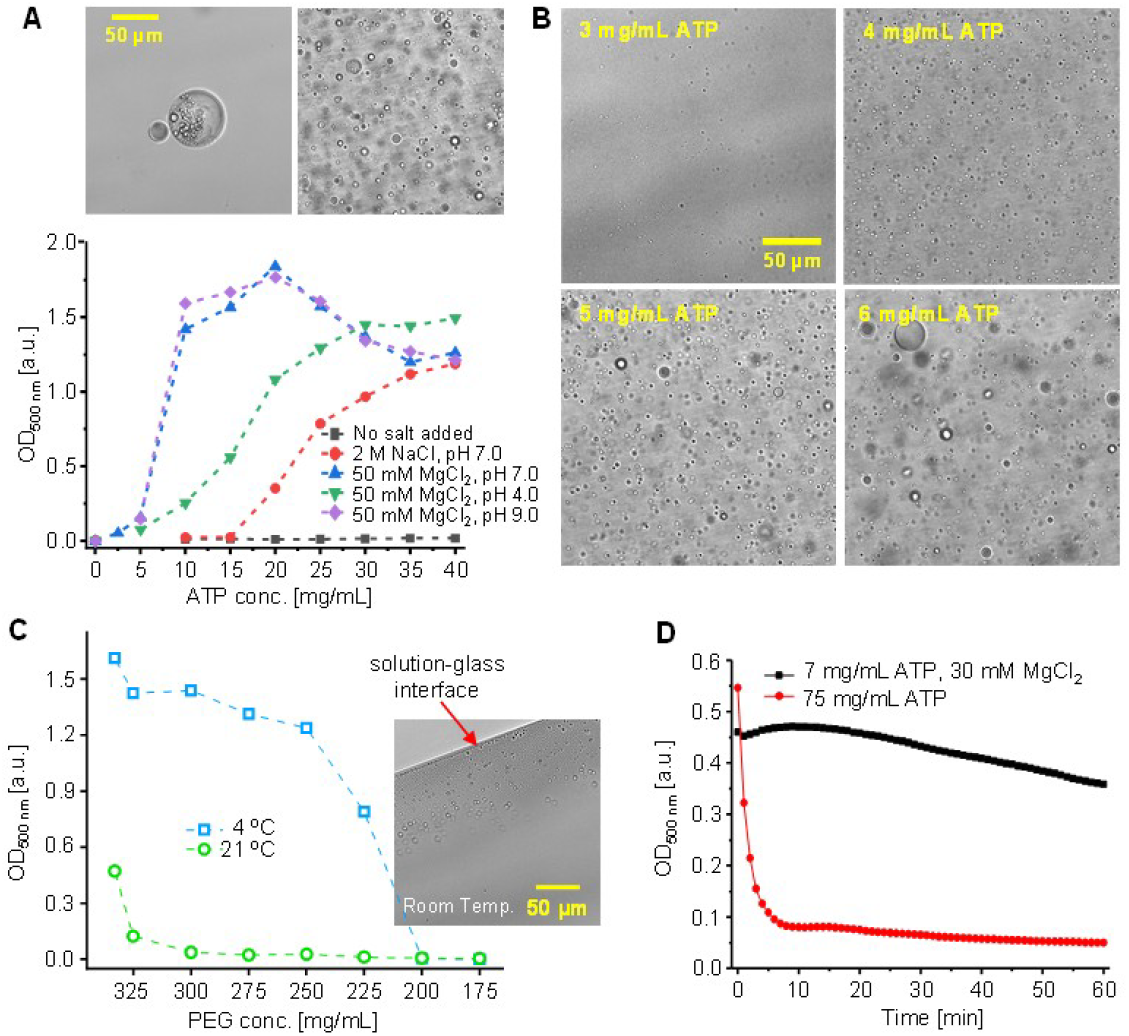
Moderate concentrations of Mg^2+^ ions shift the LLPS transition to the physiological range of ATP concentrations. Course of LLPS-triggered changes in optical density at 21 °C with increasing ATP concentration in the absence and presence of 2 M NaCl, and in the presence of 50 mM MgCl_2_ at pH 4, 7, and 9; the PEG-3350 conc. being 333 mg/mL (A). The left and right microscopy images shown in the inset correspond to samples containing initially subcritical ATP concentrations (2.5 and 10 mg/mL, respectively) with 50 mM MgCl_2_, pH 7, and PEG-3350 at 333 mg/mL Brightfield images of droplets stabilized by 30 mM MgCl_2_ and 333 mg/mL PEG-3350, pH 7, of ATP at various concentrations in the close-to-cellular range (B). Optical density dependency on PEG concentration in the Mg^2+^-containing ATP samples (7 mg/mL ATP, 30 mM MgCl_2_, pH 7) at 21 °C and 4 °C (C). The inset image shows how tiny droplets are formed at the solution-glass interface even at room temperature and subcritical PEG concentration of 175 mg/mL. Tiny Mg^2+^-stabilized droplets coalesce more slowly than droplets formed at high ATP concentrations, as reflected by different rates of optical density changes in quiescently sedimenting samples (D – 7 mg/mL ATP + 30 mM MgCl_2_, compared to 75 mg/mL ATP, both in the presence of 333 mg/mL PEG-3350, pH 7).

### The effects of adenosine di- and monophosphate and different crowder types on phase separation and droplet formation

Additional experiments conducted in the course of this work have shown that both ADP and AMP (adenosine di- and monophosphate) also form droplets in the presence of concentrated PEG, although the droplet state is less stable at low pH in either case and tends to undergo liquid-solid phase transition (LSPT), i.e., crystallization (Figs. S2 and S3). We also note that all three nucleotides can form droplets cooperatively when each of them is present separately in a subcritical concentration (Fig. S4). Finally, it should be emphasized that although a change in the molecular weight of PEG has only a minor effect on LLPS in aqueous ATP solution, replacing PEG with the polysaccharide-like crowding agents Ficoll-70 or Dextran-10 render the transition no longer observable under the typical set of physicochemical conditions (Fig. S5). Hence, the pronounced surface hydrophobicity of PEG in concert with the excluded volume effect must be responsible for triggering the phase separation.

## Conclusions

In summary, we have discovered that ATP — the cell’s central metabolite — has the intrinsic capacity to undergo liquid–liquid phase separation in the absence of proteins or peptides. The amphilic nature of ATP, presenting Coulombic, hydrophobic, π-stacking and possibly also entropically favorable anion-π interactions (52), permits stability and dynamics of the ATP droplets over a wide range of pH values and ionic strengths. Under physiologically relevant crowding conditions and with millimolar Mg^2+^, this phase separation occurs at intracellular ATP concentrations. These findings elevate ATP from a mere modulator of protein condensation to an autonomous phase-separating species capable of forming its own condensates. Local increases or fluctuations in ATP and Mg^2+^ levels can therefore directly nucleate or regulate membraneless organelles, while dysregulated ATP phase behavior could contribute to pathological condensation pathways. This previously overlooked facet of ATP biochemistry opens new avenues for understanding cellular organization and the molecular origins of condensate-related diseases.

## Materials and methods

### Samples

PEG-2000 was purchased from Fluka Biochemika (Honeywell Research Chemicals), Dextran-10 from Carl Roth, Ficoll-70 from GE Healthcare Bio-Sciences AB, PEG-FITC (MW 3350) from Creative PEGWorks, the ATP determination kit from Invitrogen (Thermo Fisher Scientific). ATP, ADP, AMP, PEG-3350, PEG-8000, quinacrine, NaCl, MgCl_2_ and all other reagents were purchased from MilliporeSigma (formerly Sigma-Aldrich).

Samples were prepared by suspending and dissolving weighted amounts of ATP (or ADP/AMP) and crowder in water at approximately 90 % of the targeted total volume, followed by pH-adjustment of stirred liquids with NaOH or HCl to the desired values followed by an addition of water to the total targeted volume and confirmation of the final pH value. In the case of NaCl or MgCl_2_-containing samples, the procedure was modified so that the specific salt was present in the sample at the desired concentration.

### Optical microscopy

Light microscopy images were recorded on an Eclipse TE2000-U microscope (Nikon Inc.) equipped with a Nikon Plan Fluor 10x objective (0.45, WD 4.0) and additionally a X-Lite™-120 fluorescence Illumination System EXFO and a FITC-HQ filter (excitation 480/40, emission 527/30) for the PEG-FITC and quinacrine fluorescence mode.

### Optical density (turbidity) measurements

Optical density was measured at 500 nm using a UV-1800 Shimadzu UV Spectrophotometer in a 3 mm pathlength cuvette. Unless otherwise indicated, measurements were carried out at 21 °C. Samples were vortexed directly before the measurement (also in the case of measurements related to cooling and testing the effect of elevated temperatures).

### Luciferase-based ATP assay

To estimate the relative ATP concentration in the top and bottom phases as well as in their mix, an ATP assay based on the bioluminescent reaction of ATP with luciferin catalyzed by luciferase was used. Measurements were performed on a CLARIOstar® plate reader (BMG LABTECH) using a 96-well black microplate, recording luminescence at 560 nm for samples diluted 1:9100 and at a temperature of 28 °C.

### Circular dichroism (CD) spectroscopic measurements

For the CD measurements, the dense droplet phase on the ATP-PEG system (obtained at the specified pH) system was centrifuged and placed in a demountable quartz cuvette of 0.01 mm optical path. The CD spectrum, corrected for the buffer signal, was acquired at 25 °C by accumulation of 20 independent spectra (at 200 nm/s scanning rate) on a J-815 S spectropolarimeter from Jasco Corp. (Tokyo, Japan).

## Funding

The authors acknowledge funding from the Deutsche Forschungsgemeinschaft (DFG, German Research Foundation) under Germany’s Excellence Strategy – EXC 2033 – project number 390677874-RESOLV. WD acknowledges support from the University of Warsaw’s IDUB program (BOB-661-501/2025).

## Author contributions

R.D. performed the experiments; W.D. and R.W. managed the project; all authors wrote the manuscript.

## Competing interests

Authors declare that they have no competing interests.

## Data and material availability

All data are available in the main text or the supplementary materials.

## SUPPLEMENTARY MATERIALS

Figs. S1-S5: Stability of the two-phase ATP-PEG system at room temperature; LLPS and LSPT in the ADP-PEG system; LLPS and LSPT in the AMP-PEG system; LLPS in mixed ATP+ADP+AMP-PEG system; ATP droplets: PEG-3350 vs. other macromolecular crowders.

## Notes

### Competing Interest Statement

The authors have declared no competing interest.

