## supplementary information for "Protein-Independent Liquid–Liquid Phase Separation of Adenosine Triphosphate under Crowded Conditions"

for

**CONTENTS**

1. Stability of the two-phase ATP-PEG system at room temperature
2. LLPS and LSPT in the ADP-PEG system
3. LLPS and LSPT in the AMP-PEG system
4. LLPS in mixed ATP+ADP+AMP-PEG system
5. ATP droplets: PEG-3350 vs. other macromolecular crowders

#### 1. Stability of the two-phase ATP-PEG system at room temperature

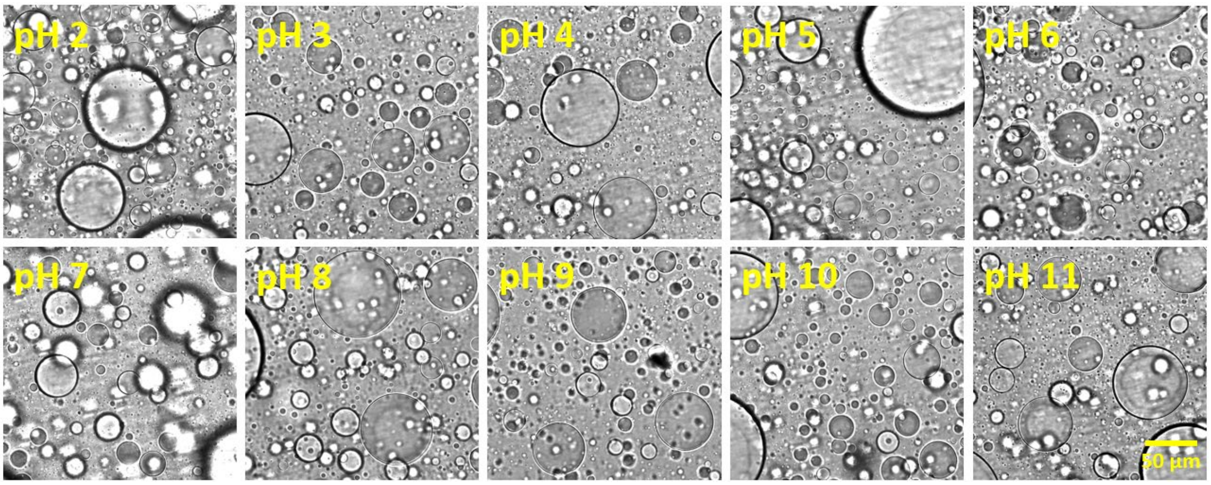


**Fig. S1.** Brightfield microscopic images of ATP-PEG-3350 droplets after 7 days at room temperature at different pH values (samples shaken before the measurement). Samples: 75 mg/mL ATP, 333 mg/mL PEG (average molecular weight 3350), $H_2O$; pH as specified in the figure; room temperature (including during aging).

### 2. LLPS and LSPT in the ADP-PEG system

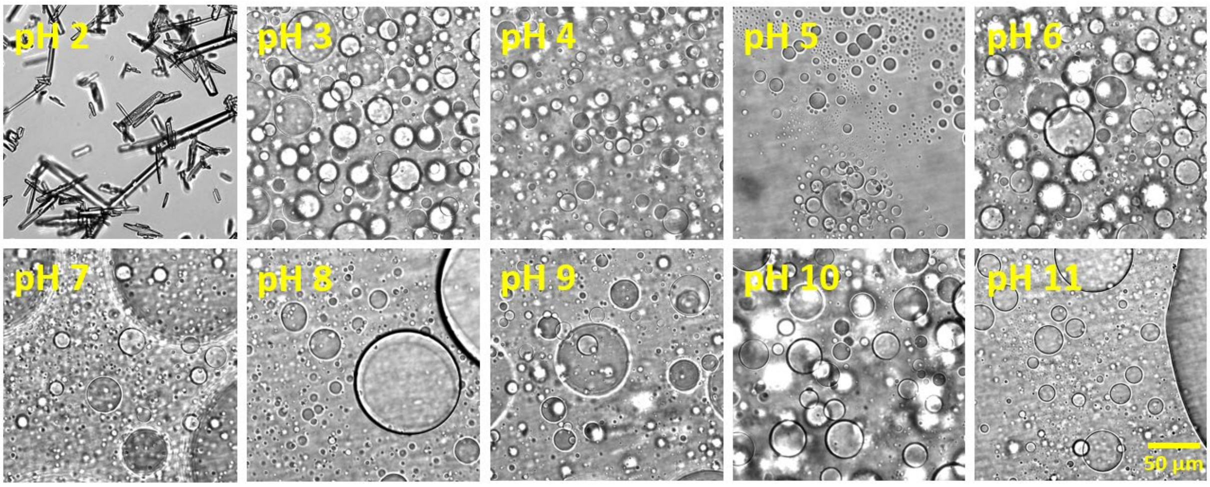


**Fig. S2.** Brightfield microscopic images of the ADP-PEG-3350 system: droplets also form, but at very low pH, the system reveals a stronger tendency to form crystals. Samples: 75 mg/mL ADP, 333 mg/mL PEG (average molecular weight 3350), $H_2O$; pH as specified in the figure; room temperature.

### 3. LLPS and LSPT in the AMP-PEG system

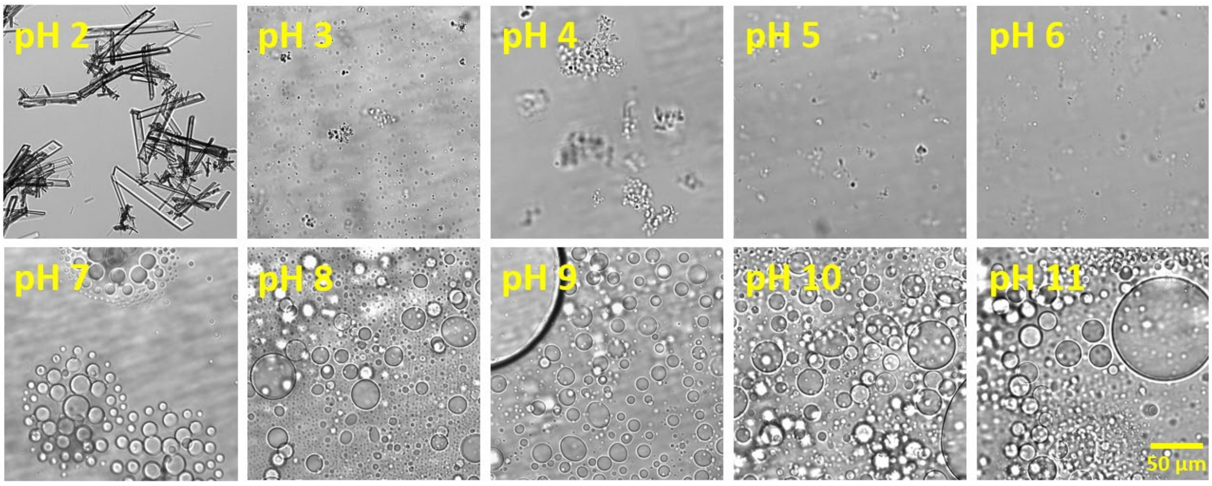


**Fig. S3.** Brightfield microscopy images of the AMP-PEG-3350 system: crystals or solid aggregates dominate at low pH, whereas droplets prevail at pH 7 and above. Samples: 75 mg/mL AMP, 333 mg/mL PEG (average molecular weight 3350), $H_2O$; pH as specified in the figure; room temperature; equipment: Eclipse TE2000-U microscope.

### 4. LLPS in the mixed ATP+ADP+AMP-PEG system

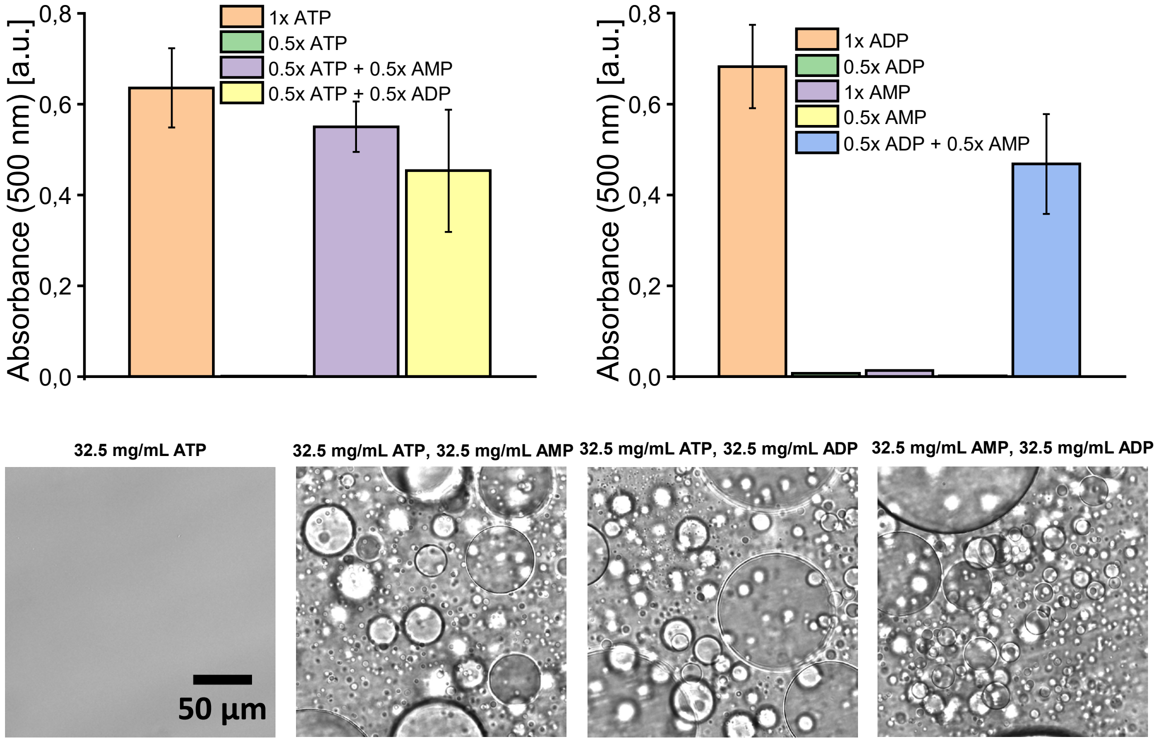


**Fig. S4.** ATP, ADP, and AMP in PEG at concentrations below their individual droplet-permissive concentration thresholds form droplets when mixed together. Comment: ATP, ADP, and AMP in PEG do not form light-scattering droplets when used at a concentration of 32.5 mg/mL (0.5× in the graphs). However, if samples contain simultaneously ATP and ADP (or ATP/AMP, ADP/AMP), each at 32.5 mg/mL, strong absorbance and numerous droplets are observed.

### 5. ATP droplets: PEG-3350 vs. other macromolecular crowders

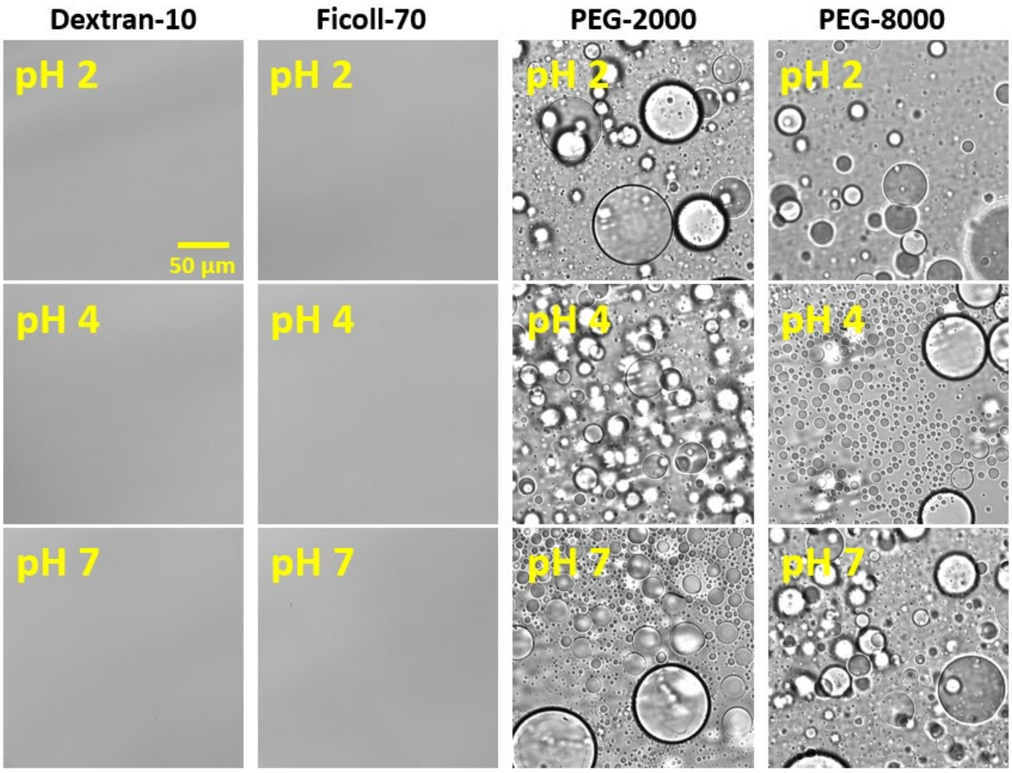


**Fig. S5.** Brightfield microscopy images of ATP in the presence of alternative macromolecular crowders. Droplets are still present in the ATP–PEG system when PEG-3350 is replaced by PEG-2000 or PEG-8000, but are absent if PEG is replaced with Dextran-10 or Ficoll-70. Comment: PEG, due to its amphiphilic character, can sequester water and enhance hydrophobic interactions. Dextran and Ficoll do not have this property. Samples: 75 mg/mL ATP, 333 mg/mL molecular crowder (Dextran-10, Ficoll-70, PEG-2000, or PEG-8000), $H_2O$; pH as specified in the figure; room temperature; equipment: Eclipse TE2000-U microscope.
